# Multilevel proteomic profiling of colorectal adenocarcinoma cell differentiation to characterize an intestinal epithelial model

**DOI:** 10.1101/2024.02.19.581096

**Authors:** Emily EF Fekete, Angela Wang, Marybeth Creskey, Sarah E Cummings, Jessie R Lavoie, Zhibin Ning, Jianjun Li, Daniel Figeys, Rui Chen, Xu Zhang

## Abstract

Emergent advancements on the intestinal microbiome for human health and disease treatment necessitates well-defined intestinal cellular models to study and rapidly assess host, microbiome, and drug interactions. This study characterized molecular alterations during Caco-2 cell differentiation, an epithelial intestinal model, using quantitative multi-omic approaches. We demonstrated that both spontaneous and medium-induced cellular differentiations displayed similar protein and pathway changes, including the down-regulation of proteins related to translation and proliferation, and up-regulation of proteins related to cell adhesion, molecule binding and metabolic pathways. Acetyl-proteomics revealed decreased histone acetylation and increased acetylation in proteins associated with mitochondria functions in differentiated cells. Butyrate-containing differentiation medium accelerates differentiation, with earlier up-regulation of proteins related to differentiation and host-microbiome interactions. These results emphasize the controlled progression of Caco-2 differentiation toward a specialized intestinal epithelial-like cell. This further enhances their characterization, establishing their suitability for facilitating the effective evaluation of risk and quality in microbiome-directed therapeutics.

## Introduction

The gastrointestinal (GI) tract is an important collection of organs in the human body, which is involved in various biological processes, including nutrition and immune development ^1^. Diseases originated from GI tract, such as colorectal cancer and inflammatory bowel diseases (IBD), are among the most common diseases in modern society ^1,2^. The GI tract is also the home of diverse human symbiotic microbial communities, namely the human microbiome ^1^. The gut microbiome in particular has gained considerable attention in recent years due to its profound impact on human health and disease ^1,2^. Increasingly, causative roles of gut microbiota in diseases have been identified, leading to the intensive development and application of microbiota-directed therapeutics, such as the use of fecal microbiota transplantation (FMT) for preventing recurrent *C. difficile* infection ^3–5^. The establishment of appropriate *in vitro* models that accurately recapitulate host-microbiome interactions is crucial for advancing our understanding of the complex interplay between the human body and its resident microbial communities, as well as the efficient risk and quality assessment of therapeutics targeting or derived from microbiomes ^3^.

The intestinal epithelial barrier plays a vital role in host-microbiome interactions ^1^. Caco-2 cells, derived from human colon adenocarcinoma, have been widely employed as a prominent epithelial model for drug and intestinal disease studies due to their ability to mimic various aspects of intestinal physiology and provide insights into pathology of the intestinal epithelium ^6,7^. Caco-2 cells have demonstrated the ability to undergo differentiation into epithelium-like cells through both spontaneous and butyrate-induced processes ^6,8–12^. Spontaneous differentiation of Caco-2 cells is primarily driven by the intrinsic program of the cells themselves. Once reaching confluence and establishing contact inhibition, cell-to-cell contact and communication generates a physical cue for the cells to differentiate ^6,9,12^. The addition of butyrate, a metabolite derived from the microbial fermentation of dietary fiber, has been shown to promote intestinal epithelial cell differentiation ^8,12^. In both cases, there is a progression away from a proliferative state towards a more differentiated and specialized phenotype over the course of one to a few weeks depending on differentiation methods used ^6,8,9,12^.

To enable the application of the Caco-2 cellular model in microbiome drug development and host-microbiome interaction study, there is a need to revisit and comprehensively characterize the molecular processes at play during differentiation. Previous studies have examined transcriptomic, proteomic and glycomic alterations that Caco-2 cells undergo upon either spontaneous or butyrate-induced differentiation ^9,10,12^. In this study, we took advantage of tandem mass tag (TMT) multiplexing and enrichment strategies to further these previous efforts for in-depth, proteome-wide and multilevel characterizations of the Caco-2 differentiation over a time course, including quantitative proteomics, lysine acetylproteomics and glycoproteomics ^13^. We compared three commonly used differentiation protocols including both spontaneous (using serum-free and serum containing media) and differentiation medium (DM)-induced differentiation. The multilevel –omic characterizations of multiple Caco-2 differentiation protocols performed in this study provide greater insights into the application of the Caco-2 cellular model for the study of host-microbiome interactions and the assessment of microbiome-directed therapeutics.

## Results

To characterize the Caco-2 cell differentiation, confluent cells were cultured using either regular Dulbecco’s Modified Eagle Medium (DMEM) growth media with serum (DFBS), without serum (SFM), or a commercial enterocyte differentiation media (DM, contains butyrate) for 21 days (Fig. 1A). It is reported that DM can induce full cellular differentiation in 5-7 days ^8^. Accordingly, we found that at day 7, DM group cells formed dome structures, hallmark of Caco-2 cellular differentiation, but not yet for DFBS and SFM groups (Supplementary Fig. 1A-C). To examine and compare the molecular alterations at the proteome level, we collected samples throughout the course of differentiation at day 1, day 3, day 7, day 14, and day 21 (Fig. 1A). The harvested cells were then lysed, digested, labelled with isobaric TMT tags followed by either total proteome measurement or further enrichment by immunoaffinity approach for lysine acetyl-proteomics and hydrophilic interaction liquid interaction chromatography (HILIC) for glycoproteomics (Fig. 1B).

**Figure 1.**
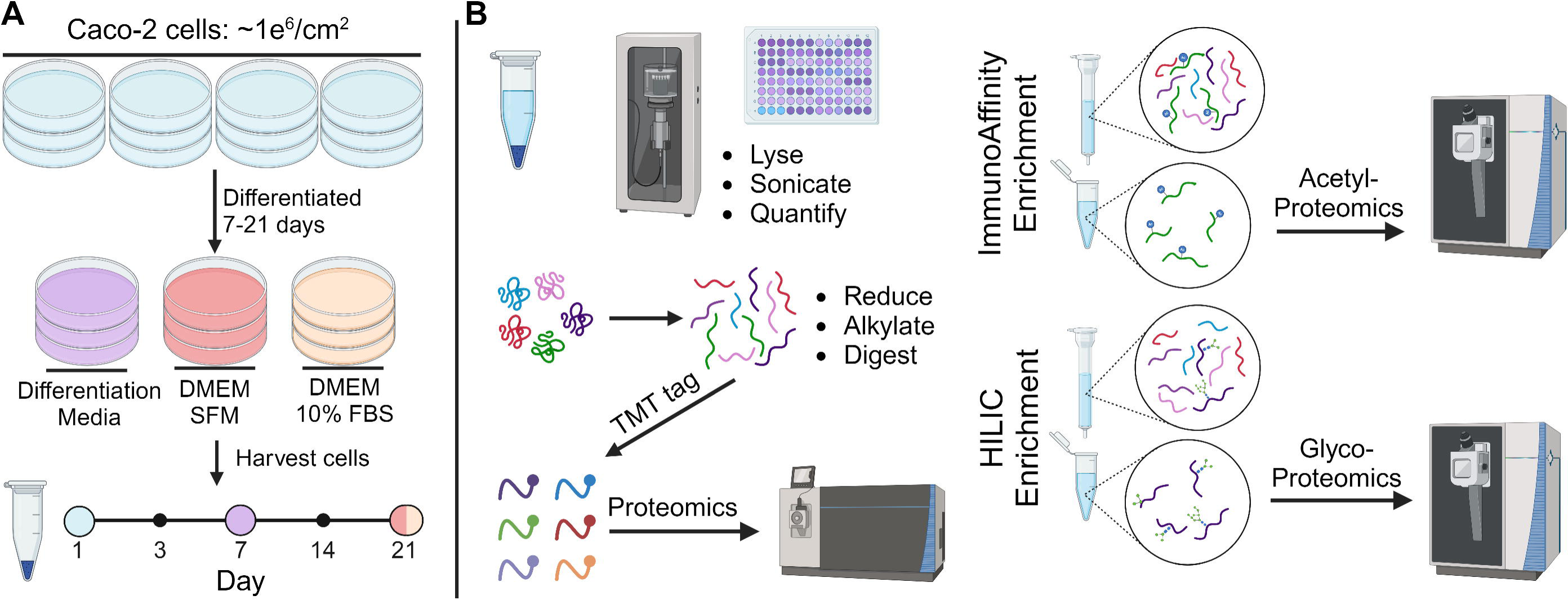
Graphic scheme depicting the cell culture and multi-omics workflow of this study. Caco-2 cells cultured in different media over 21 days with sample collection at different time points (A). Sample processing workflow used to prepare peptide samples for mass spectrometry (MS) analysis (B).

### Quantitative proteomics demonstrates differentiation of Caco-2 cells

For multiplexed quantitative proteomics, a total of 54 samples (6 replicates for undifferentiated cells on day 1, and 4 replicates for all other groups) were randomized and assigned into six TMT11plex experiments (Supplementary Table 1) ^13^. To allow for cross-TMT experiment normalization, channels 1 and 11 of each TMT experiment were used to label a mix reference sample. Each of the TMT experiment mixtures were then fractionated into 8 fractions and analyzed by mass spectrometry (MS). The resulting raw MS files were then processed with MaxQuant, which identified 45, 555 peptides and 5038 protein groups. MSstatsTMT was then used for summarization and normalization, which resulted in the quantification of 4962 protein groups in total, and 3337 were quantified in all samples.

Principal Component Analysis (PCA) of the quantified protein groups in all samples was performed to examine the clustering patterns of undifferentiated Caco-2 cells compared to cells undergoing spontaneous and DM-induced differentiation over time (Fig. 2A). The PCA score plot revealed distinct clustering of the different groups, indicating significant differences in their protein expression profiles. Undifferentiated and differentiated Caco-2 cells, as well as Caco-2 cells which underwent different differentiation protocols, over time clustered separately from each other. This observation suggests distinct proteomic changes in differentiated cells compared to undifferentiated cells, as well as distinct proteomic changes between spontaneous and DM-induced differentiated Caco-2 cells.

**Figure 2.**
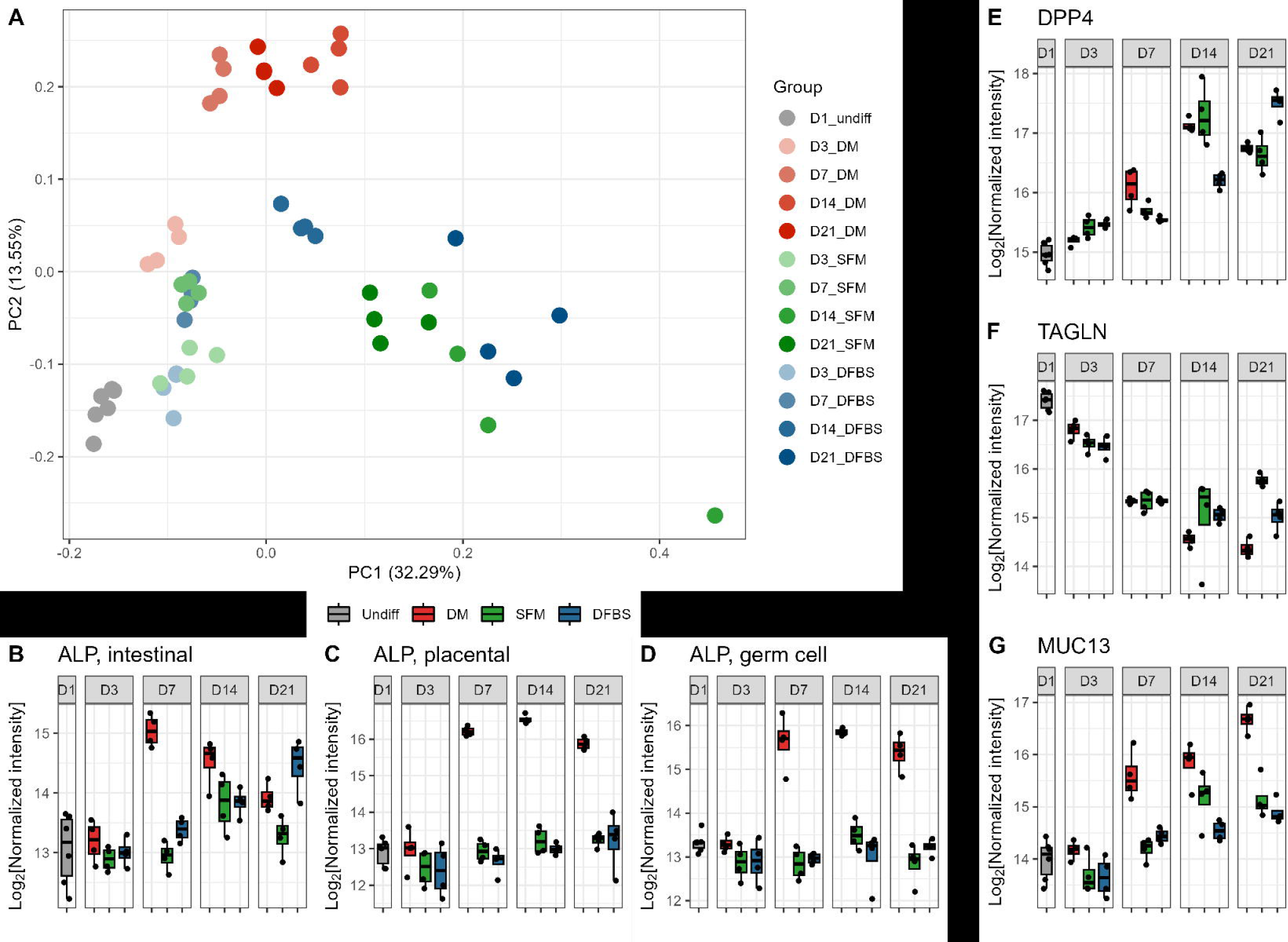
Proteomic overview and established markers of Caco-2 differentiation from carcinoma into intestinal epithelial-like cells. PCA score plot of quantified proteins across all samples (A). Log_2_(Normalized Intensity) of alkaline phosphatase (ALP) intestinal (B), placental (C), and germ cell (D) isoforms over time. Log_2_(Normalized Intensity) of other selected differentiation markers; DPP4 (E), TAGLN (F), MUC13 (G).

We then evaluated the expression patterns of known biomarkers for intestinal epithelium cell differentiation (Fig. 2B-G). Alkaline phosphatase (ALP) activity has long served as a marker of Caco-2 cell differentiation ^14,15^. It is expected that protein expression levels would positively correlate with alkaline phosphatase activity. In the DM group, a significant increase in intestinal type ALP abundance was observed as early as day 7 (adj. p-value 9.8E-13) (Fig. 2B). The DFBS group displayed a more gradual increase in ALP abundance over time, taking until day 21 to have a significant rise in ALP abundance (adj. p-value 5.8E-09)(Fig. 2B). Interestingly, we also noticed a significant increase in placental (adj. p-value 7.0E-17) and germ cell type (adj. p-value 1.1E-11) ALP abundance within the DM group that has been previously reported in studies exploring other butyrate-treated colonic adenocarcinoma cell lines (Fig. 2C-D) ^16^. This observation suggests a potentially broad effect of butyrate on various ALP isoforms.

Additional established biomarkers for Caco-2 differentiation into intestinal epithelium-like cells observed in this study include an increase in dipeptidyl peptidase 4 (DPP4) ^17,18^, mucin 13 (MUC13) ^19^, thiosulfate sulfurtransferase (TST) ^9^, 3-hydroxybutyrate dehydrogenase 2 (BDH2) ^9^, and a decrease in transgelin (TAGLN) expression ^9^ (Fig. 2E-G & Supplementary Table 2). Villin-1 and villin-2 are both cytoskeletal proteins that are highly expressed in the brush border of intestinal epithelial cells, playing roles in maintaining the structural integrity and functionality of the microvilli, which increase the surface area for intestinal nutrient absorption and barrier function ^20,21^. We observed up regulation of villin proteins in the DM group at day 7, but not in DFBS and SFM groups (Supplementary Fig. 2G-H).

### Multivariate statistics identified differentially abundant proteins in differentiated Caco-2 cells

To identify proteins that were altered during Caco-2 cell differentiation, we performed a Partial least squares-discriminant analysis (PLS-DA) for all quantified proteins in undifferentiated (D1_undiff) and differentiated cell groups (D7_DM, D21_DFBS, D21_SFM). Successful group discrimination model was established with a goodness of prediction (Q2) of 0.89 and R2 of 0.99. By using a variable importance projection (VIP) cut-off of 1, a total of 1008 proteins were identified as differentially abundant proteins (Supplementary Table 2). Hierarchical clustering of the expression changes of all the VIP proteins indicated that 508 proteins were upregulated and 500 proteins were downregulated in differentiated cells compared to undifferentiated samples (Fig. 3A). Interestingly, 135 up regulated proteins in differentiated cells have a general trend of a higher extent of up regulation in DM than SFM and DFBS groups, despite less differentiation time used for DM (7 days compared to 21 days in the other two groups) (Fig. 3A).

**Figure 3.**
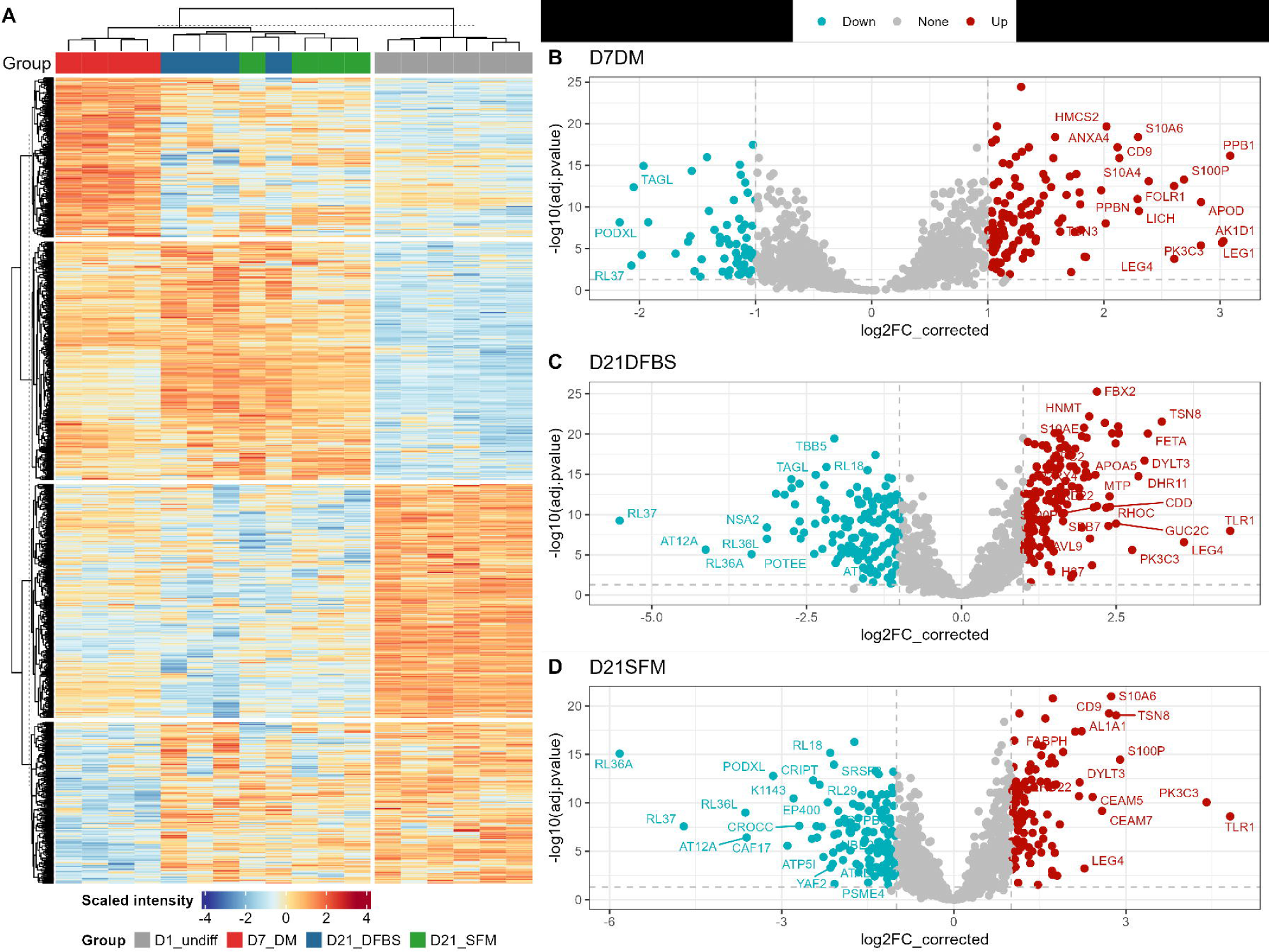
Differentially abundant protein expression of key Caco-2 cell differentiation groups. Heatmap and hierarchical clustering of samples using differentially abundant VIP ≥ 1 proteins (A). Volcano plots of Log2FC of all VIP ≥ 1 proteins, highlighting those with a fold change ≥ 2 and adjusted p-value ≤ 0.05 comparing undifferentiated day 1 Caco-2 cells to cells grown in DM for 7 days (B), DFBS for 21 days (C), and SFM for 21 days (D).

Among the up-regulated proteins in differentiated Caco-2 cells when compared to undifferentiated cells and ranked highest in the PLSDA VIP analysis is phosphatidylinositol 3-kinase catalytic subunit type 3 (PIK3C3, VIP = 7.38) which was significantly changed in all groups (Fig. 3A-C, Supplementary Fig. 2L & Supplementary Table 2). PK3C3 is recognized to be involved in multiple membrane trafficking pathways, suggesting extensive membrane re-structurization during Caco-2 differentiation. CD9 (VIP = 4.97), a cell surface glycoprotein and well-known exosome biomarker, was also among the highest VIP ranking significantly upregulated proteins, and showed a steady increasing trend during the entire 21 days culturing in all three differentiation protocols (Supplementary Table 2 & Supplementary Fig. 2K). We also observed that galectin-1 (VIP = 6.24), aldo-keto reductase family 1 member D1 (AK1D1, VIP = 5.99) and folate receptor alpha (FOLR1, VIP = 5.50) were among the proteins with highest VIP values, but were only or more extensively up-regulated in DM group (Supplementary Table 2 & Supplementary Fig. 2I-J).

We identified four S100 family proteins with VIP values more than 1, including S100A6 (VIP = 6.31), S100A4 (VIP = 5.25), S100P (VIP = 6.56), and S100A11 (VIP = 3.55). All of the quantified S100 family proteins showed upregulation with differentiation across both DM-induced and spontaneously differentiated groups (Fig. 4A-I &SupTable2). The expression levels of S100A4 and S100A11 significantly increased in DM group starting from day 7 when compared to other groups. The S100 protein family is a group of small, calcium-binding proteins that play diverse roles in cellular functions ^22^. In addition to S100 family proteins, annexins are another family of calcium-dependent proteins that are involved in diverse cellular processes, including membrane dynamics, vesicle trafficking, and signal transduction and can form protein complexes with S100 proteins in certain cellular contexts ^23^. Accordingly, we also found that many annexins, in particular annexin A6, were increased in DM induced differentiated cells only and the increase starts as early as day 3 of induction (Supplementary Fig. 2A-F).

**Figure 4.**
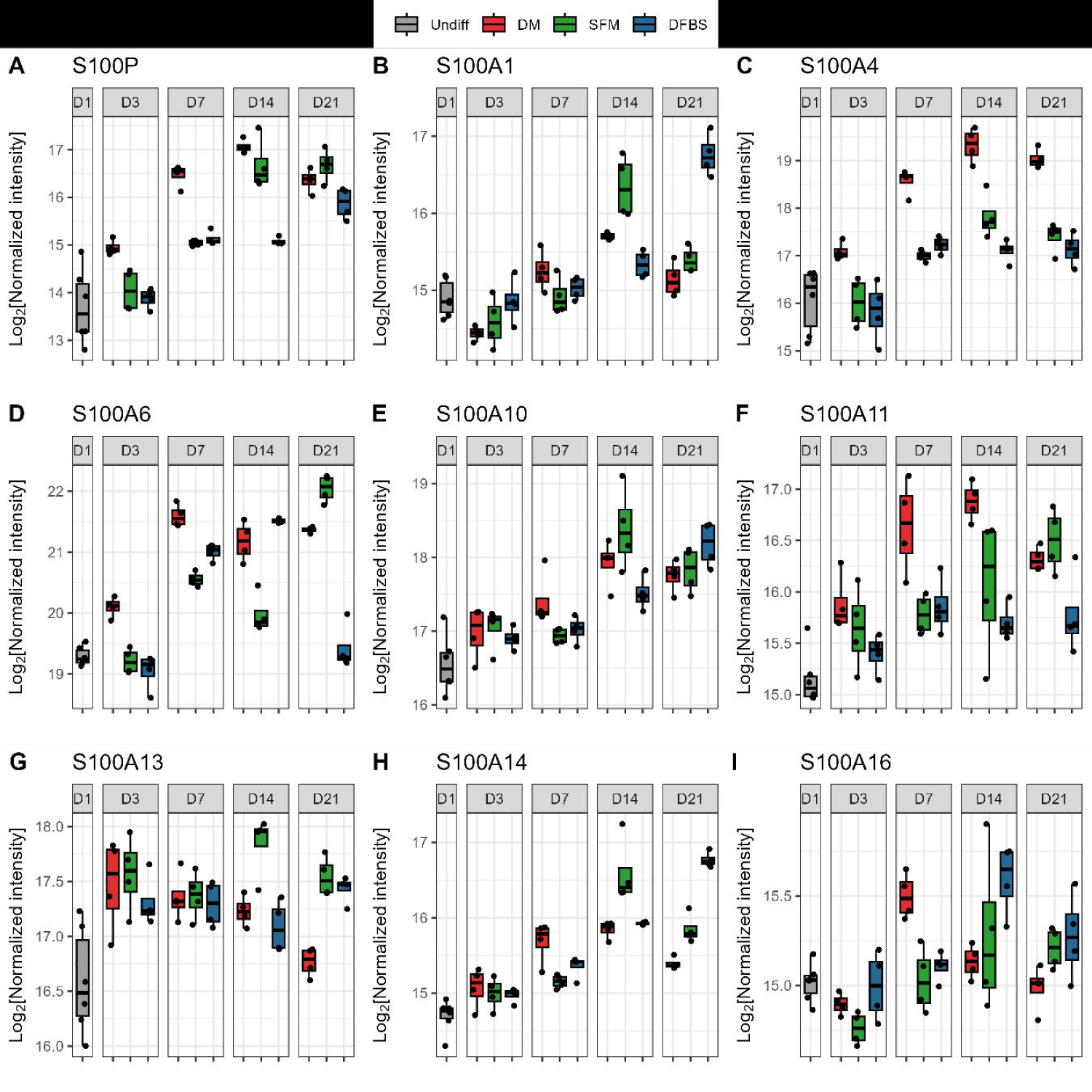
Differentially abundant protein expression of key Caco-2 cell differentiation groups. Log_2_(Normalized Intensity) of S100 protein family members; S100P(A), S100A1(B), S100A4(C), S100A6(D), S100A10(E), S100A11(F), S100A13(G), S100A14(H), and S100A16(I) over time.

In agreement with the retarded cell proliferation in differentiated cells, we found that the most significantly down-regulated proteins include ubiquitin-conjugating enzyme E2 T (VIP = 4.74), large ribosomal subunit protein eL37 (RPL37, VIP = 5.28), eL42 (RPL36A, VIP = 4.80), and DNA-directed RNA polymerase I subunit RPA34 (CD3EAP, VIP = 2.69) (Supplementary Fig. 2 M-P & Supplementary Table 2). We also found a significant decrease in the expression level of podocalyxin (PODXL, VIP = 5.14), an anti-adhesive transmembrane glycoprotein, during Caco-2 cellular differentiation in all three protocols. Over-expression of PODXL has been reported to inhibit cell-cell interaction and be indicative of poor prognosis in colorectal cancer ^24^. The decrease of PODXL is therefore in agreement with the fact that differentiated Caco-2 cells have well established tight junctions.

### Functional and pathway alterations in differentiated Caco-2 cells

To better explore the functional and pathway changes during differentiation, we then performed functional enrichment analysis for the up- or down- regulated proteins in differentiated Caco-2 cells using STRING database (Fig. 5).

**Figure 5.**
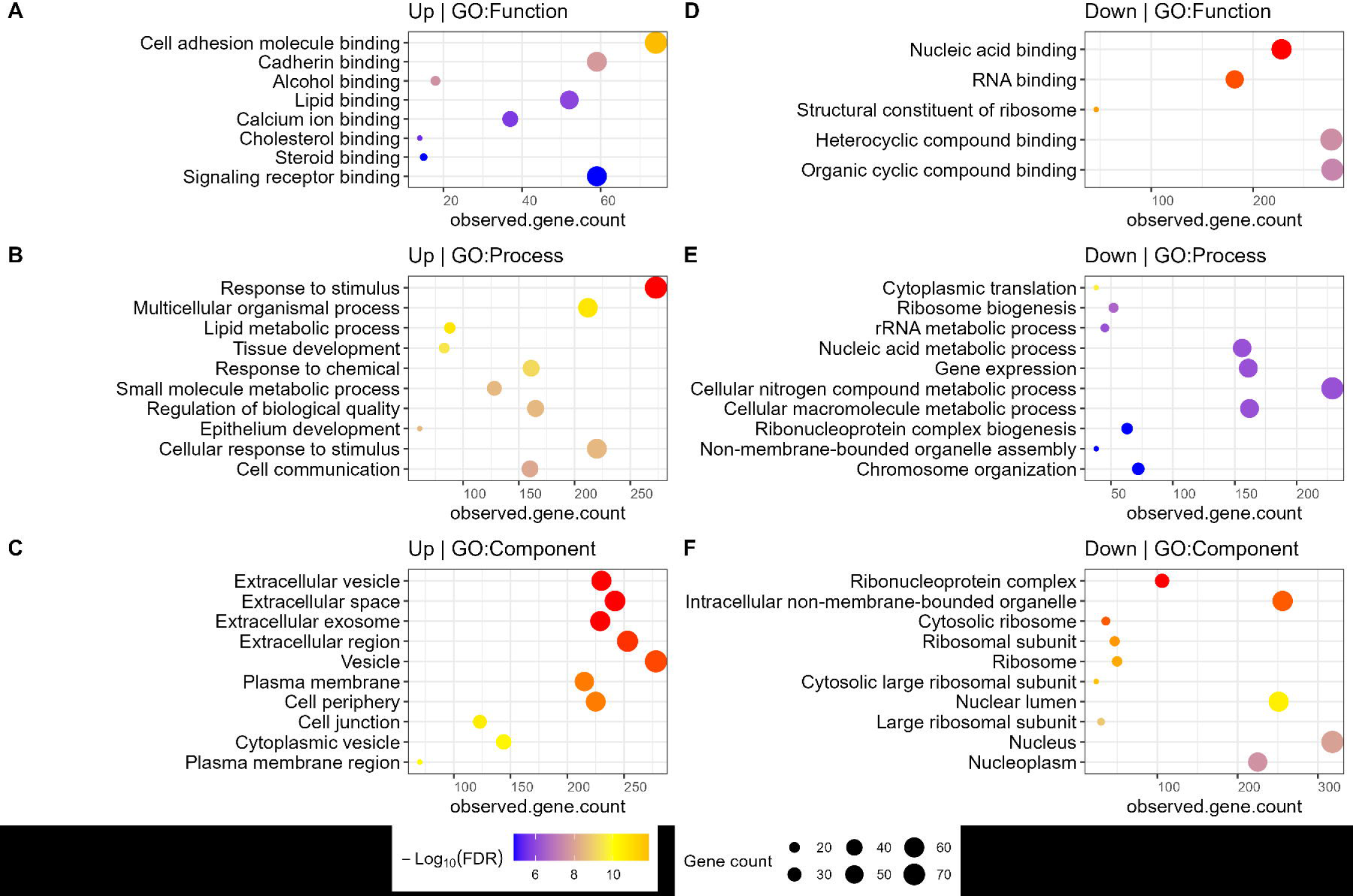
Clustering and functional enrichment analysis of differentially abundant proteins in differentiated cells compared to undifferentiated Caco-2 cells. Gene Ontology enrichment analysis using STRING for PLSDA VIP ≥ 1 proteins that were up-regulated (A-C) and down-regulated (D-F), respectively, in differentiated groups (D7_DM, D21_DFBS or D21_SFM).

The up-regulated proteins in the differentiated groups were enriched in functions related to cell adhesion molecule binding, cadherin binding, calcium ion binding and other metabolite/small molecule/protein/lipid binding activities (Fig. 5A). Biological processes related to the responses to stimuli and chemicals, lipid metabolism, cell communication, as well as epithelial development were significantly enriched in the up-regulated proteins in differentiated Caco-2 cells (Fig. 5B). The up-regulated proteins were also significantly enriched in cellular components, including extracellular vesicles, exosomes, plasma membranes, and cell junctions (Fig. 5C). These observations indicate a shift in the cellular activity as it differentiates, including an increase in the formation of junctions and connections between cells, as well as the development of a more epithelial phenotype, which align with known cell-changes post-differentiation. A subset of the up-regulated proteins in differentiated cells displayed a higher extent of upregulation within the DM group (Fig. 3A). Functional enrichment analysis showed that these proteins were enriched in functions and pathways related to cell adhesion, epithelial differentiation, with a focus of proteins executing functions in extracellular vesicles, cell junctions, and around the cell membrane (Supplementary Fig. 3A-C).

The down-regulated proteins in the differentiated groups were enriched in functions mainly involved in nucleotide binding, which would align with a decrease in cell replication (Fig. 5D). In the differentiated groups there was a decrease of processes related to translation, ribosome biogenesis and nucleic acid metabolism (Fig. 5E). From the cellular components enrichment analysis the down-regulated proteins were mainly enriched in the cytoplasmic space, nucleus, and intracellular organelles (Fig. 5F). These pathway level findings align with a progression away from cell division during differentiation. Overall, these functional and pathway level observations aligned with a phenotype change from replicating carcinoma cells to a more specialized, interconnected, epithelial-cell phenotype.

### Glycoproteomic alterations during Caco-2 differentiation

Total proteome analysis demonstrated significant alterations of cell-cell interactions which may involve extensive membrane protein changes. In order to deepen the analysis of these cell surface and highly glycosylated membrane proteins, we utilized HILIC to enrich glycopeptides prior to LC-MSMS analysis. Overall, 1070 N-glycosylated sites were identified in this study, 731 of which were quantified in ≥50% of samples. The PCA score plot of the expression levels of quantified N-glycosylated protein sites matched clustering patterns seen in the proteomics analysis, showing over time distinct clustering of the different groups (Supplementary Fig. 4A).

This suggests that both spontaneous and DM-induced differentiation induced changes of N-glycoprotein expression at the cell surface compared to undifferentiated cells. The findings in this study are also in agreement and supplement the observations in a recent integrated glycomic and proteomic study, which demonstrated significant glycan structural alterations in butyrate-stimulated Caco-2 cell differentiations ^12^.

A PLS-DA analysis for key groups using quantified *N*-glycosylation site data showed similar clustering results to that of the proteomics dataset. Of 165 differentially abundant *N*-glycosylated sites identified with a VIP threshold over 1, 159 of them were up-regulated in differentiated Caco-2 cells (Supplementary Table 3). These differentially abundant sites were then subjected to functional enrichment analysis using STRING database (Fig. 6). As expected, the most significant enriched cellular components are related to membranes and extracellular spaces (Fig. 6). Significant molecular function enrichment was also obtained for receptor activity and cell adhesion molecule binding, in particular virus receptor and signaling receptor activities (Fig. 6A). Interestingly, in addition to the cell adhesion processes, biological processes involved in symbiotic interaction and viral entry into the host cell were also among the most significantly enriched processes (Fig. 6B), indicating the cellular adaption for host-microbiome interactions. Similar to the differential abundant proteins at proteome-level, the extracellular space and exosome are also among the most significantly enriched cellular components (Fig. 6C). STRING analysis also demonstrated extensive protein-protein interactions among the differentially abundant *N*-glycosylated proteins (Fig. 6D), suggesting close cross-talks between the pathways associated with Caco-2 differentiations. We identified several proteins with multiple sites that were upregulated in differentiated cells, such as lysosomal associated membrane protein 1 (LAMP-1, 10 sites), carcinoembryonic antigen-related cell adhesion molecule 1 (CEACAM-1, 7 sites), prosaposin (SAP, 4 sites), DPP4 (4 sites), CEACAM-5 (7 sites), and galectin 3 binding protein (LG3BP, 6 sites) (Supplementary Table 3). These findings in particular the pathway changes further demonstrated an increase and the involvement of glycosylation in regulating cell adhesion, signalling pathway, and membrane transport upon transition of Caco-2 to an intestinal-epithelial-like phenotype, providing further insights into the application of differentiated Caco-2 cells to study host-microbiome interactions.

**Figure 6.**
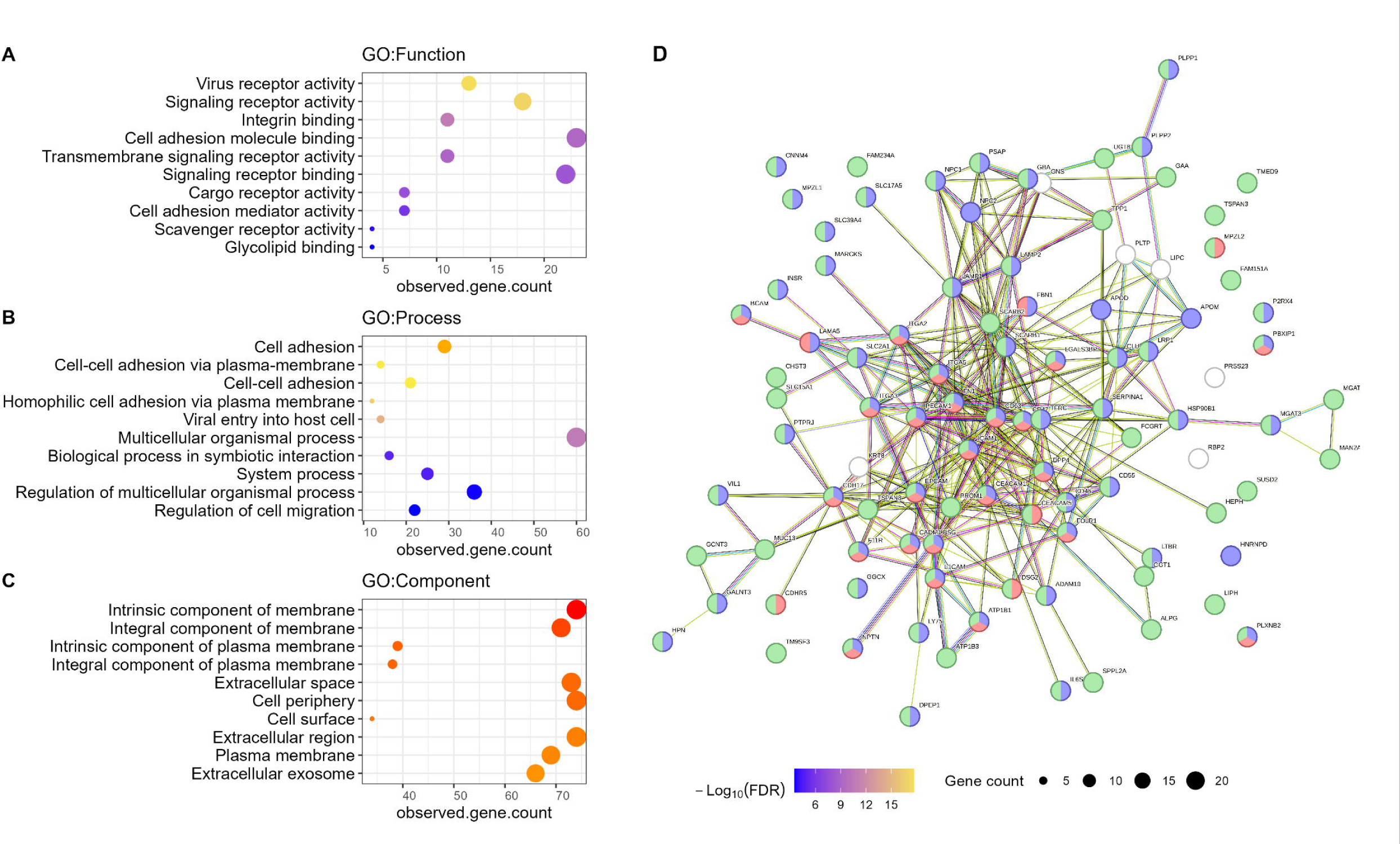
Functional enrichment and protein-protein interactions of differentially abundant N-glycosylated proteins in differentiated cells. Gene Ontology enrichment for biological functions (A), processes (B), and cellular components (C) were performed using STRING with PLSDA VIP ≥ 1 proteins upregulated in differentiated Caco-2 cells. Protein-protein interaction map were generated using STRING (D) highlighted enrichment terms are; cell adhesion (GO: 0007155, Red), response to stimulus (GO:0050896, Blue), and membrane (GO: 0016020).

### Lysine acetylation was involved in Caco-2 cell differentiations

Protein acetylation plays important roles in regulating translation, transcription, metabolism and immunity in both eukaryotes and bacteria. Aberrant protein acetylation levels have been implicated in various cancers, such as colorectal cancer ^25,26^, and dysbiotic host-microbiome interactions in diseases, such as Crohn’s disease ^27^. To this end, we performed lysine acetyl-proteomics analysis for Caco-2 differentiation. Following identification and quantification with MaxQuant, MSstatsPTM was used to summarize the acetyl-proteomic data and to perform abundance adjustment using the total proteome data. In this study, multiple lysine acetylation (Kac) modifications on one identified modified peptide sequence were considered as an individual site (highly acetylated sites; mainly present on histone proteins). Altogether, this study identified 5994 Kac sites on 2136 proteins, with the most abundant sites on histone protein H4, H31, H2B type 1-K, as well as acyl-CoA-binding protein, chloride intracellular channel protein 1 (CLIC1), malate dehydrogenase, adenosylhomocysterinase and annexin A4 (Supplementary Table 4). A total of 3459 Kac sites were quantified in >50% samples and PCA analysis showed that the undifferentiated Caco-2 cells clustered together while the cells undergoing differentiation shift away from undifferentiated cells by time regardless of differentiation protocols, suggesting altered lysine acetylproteomes of differentiated Caco-2 cells (Supplementary Fig. 5A).

To identify differentially abundant Kac sites, we used PLS-DA for the analysis of five representative groups, including D7_DM, D21_DM, D21_SFM, and D21_DFBS, which successfully (Q2 =0.59, R2 = 0.98) identified 914 differentially abundant Kac sites with a VIP threshold of 1 (Fig. 7A & Supplementary Table4). Among the 914 Kac sites, 449 sites were up-regulated and 465 were down-regulated in differentiated Caco-2 cells compared to undifferentiated cancerous cells (Fig. 7A). Pathway enrichment analysis showed that the proteins with up-regulated Kac sites were mainly involved in mitochondria and catabolic metabolism processes, while the proteins with down-regulated Kac sites were enriched in functions related to ribosome, translation, biosynthesis and nucleic acid binding (Fig. 7B-C, Supplementary Fig. 6A-D).

**Figure 7.**
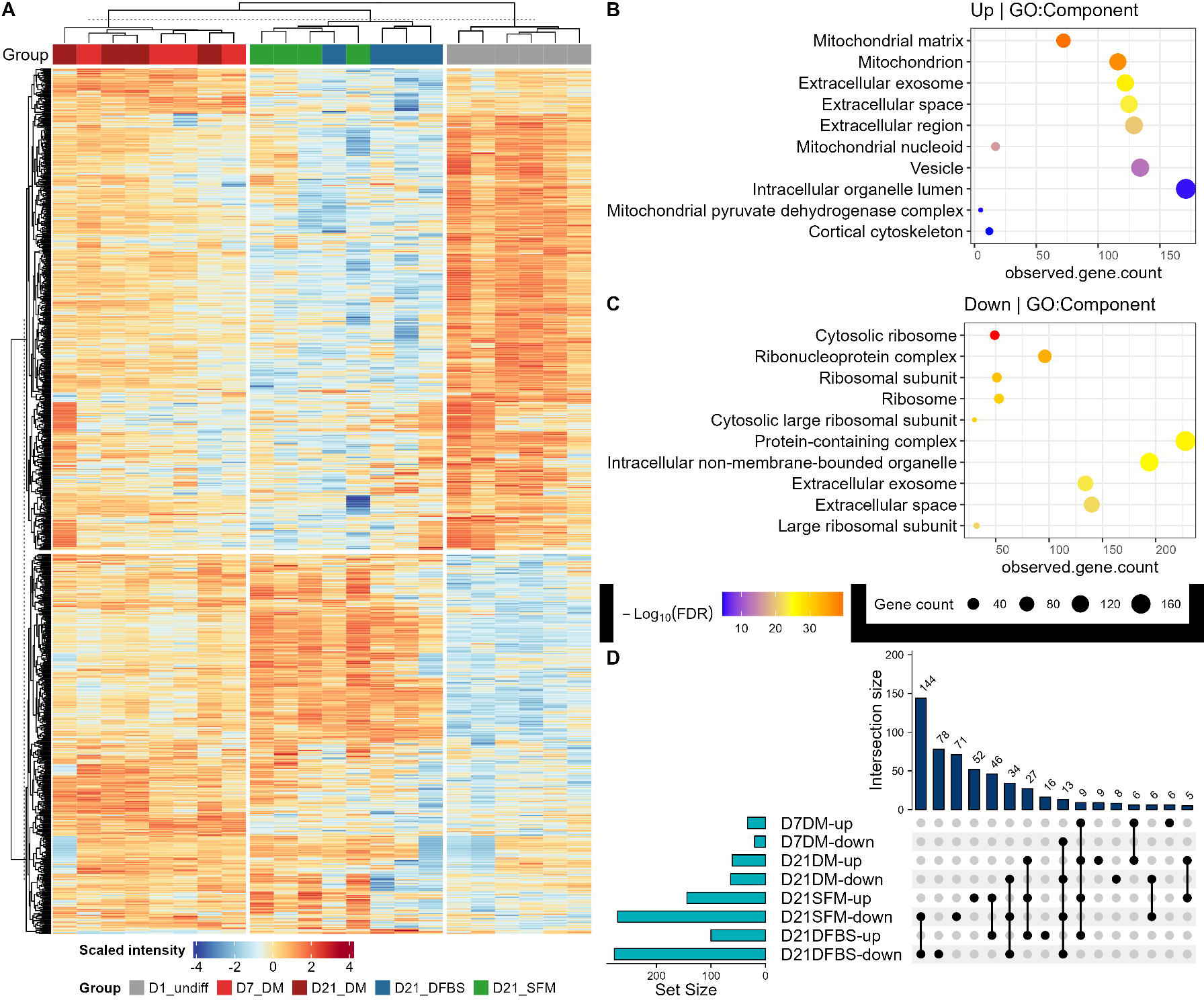
Overall trends in acetylated proteins of key differentiated Caco-2 cell groups compared to undifferentiated cells. Heatmap and clustering of PLSDA VIP ≥ 1 lysine-acetylated proteins (A). Clustering and functional enrichment analysis of differentially abundant proteins in differentiated cells compared to undifferentiated Caco-2 cells. Gene Ontology enrichment analysis using STRING for PLSDA VIP ≥ 1 of components that were up-regulated (B) and down-regulated (C) in differentiated groups (D7_DM, D21_DFBS or D21_SFM). Upset plot indicating overlap of PLSDA VIP ≥ 1, FC ≥ 2, adjusted p-value ≤ 0.05, unique lysine acetylated protein/protein sites between selected differentiated groups compared to undifferentiated Caco-2 cells (D).

Among the 914 differentially abundant Kac sites, 878 have corresponding total protein abundance quantified in proteomic dataset (Supplementary Table 4). Following adjustment using the total protein abundance, 551 Kac sites were deemed as having significantly altered acetylation levels with a threshold of fold change (FC) ≥ 2 and FDR-adjust P value ≤ 0.05. Among the 551 differentially abundant Kac sites, 53 sites (20 down, 33 up) were significantly changed in D7_DM, 125 sites (64 down, 61 up) in D21_DM, 415 sites (271 down, 144 up) in D21_SFM, and 377 sites (277 down, 100 up) in D21_DFBS groups compared to undifferentiated Caco-2 cells (Fig. 7D). The majority of the Kac sites showed consistent changing direction across all the four comparisons when overlapped (only 8 sites showed opposite directions (Fig. 7D). In agreement with the sample clustering in PCA score plot, largest overlap was found to be between D21_DFBS and D21_SFM (144 down, 46 up) (Fig. 7D). Sixty-one sites (34 down, 27 up) were found to be consistently regulated in both D21_DM, D21_DFBS and D21_SFM groups (Fig. 7D). There were 9 Kac sites being up-regulated and 13 down-regulated in all four groups, including D7_DM (Fig. 7D, Supplementary Table 4).

Histone is among the most well-known abundant and highly acetylated proteins in cells. We identified 13 differentially abundant Kac sites on 6 histone proteins (H1, H1a, H2A, H3.2, H3.3, H4) that were significantly altered (FC ≥ 2, adj. p-value < 0.05) in at least one comparison (Supplementary Table 4). The acetylation levels were decreased for most of the histone acetylation sites, except for two (K6/9 and K9/13/17) on H4 that were up-regulated in D7_DM group. The Kac site K28/37 on histone H3.2 was significantly down-regulated in all differentiation groups, except D7_DM. The acetylation level of K6/10 of histone H2A was the most significantly decreased in differentiated cells in both D21_SFM (log2FC = -4.38) and D21_DM (log2FC = -4.51) groups (Supplementary Table 4). Histone acetyltransferase p300 (p300 HAT) was also a highly acetylated protein with one triple-Kac peptide (K1542/1546/1549) identified and significantly up-regulated in all differentiation groups with a log2FC >1.55 and VIP value of 3.85 (Supplementary Table 4). HAT p300 has been previously reported to be a transcription factor coactivator and play important roles in maintaining intestinal homeostasis, cellular differentiation and prevent tumorigenesis in the intestine ^28^. The findings in this study suggest that lysine acetylation itself might be involved in regulating the function of p300, contributing to the maintenance of the intestinal cell proliferation/differentiation.

In addition to p300, the Kac sites on heat shock protein HSP90 (K255), actin-related protein 2 (ARP2, K299) and annexin A5 (K79) were also identified as up-regulated in all differentiation groups (Supplementary Table 4). We found three Kac sites (K9, K201, K246) on aldo-keto reductase family 1 member C1 protein, which were up-regulated in particular in DM groups (both day 7 and D21). On the contrary, Kac sites on ATP synthase (various on different subunits), Chloride intracellular channel protein (CLIC1, K135), and annexins A2 (K47) were more prominently upregulated in D21_DFBS and D21_SFM groups. Some of the most down-regulated Kac sites in all differentiation groups were on hepatoma-derived growth factor (HDGF, K105), alpha-enolase (K406, K335), glutathione S-transferase omega-1 (K101), vasodilator-stimulated phosphoprotein (VASP, K71), and CD9 antigen (K179) (Supplementary Table 4).

## Discussions

Differentiated monolayer Caco-2 cells are commonly used as an *in vitro* model for studies exploring intestinal absorption and reactions to various drugs or compounds. Accordingly, prior studies have explored some of the molecular changes between undifferentiated and differentiated Caco-2 cells through transcriptomics and proteomics, with most focussing in spontaneous differentiation ^6,9,12^. Microbial butyrate is known to induce rapid Caco-2 cell differentiation. Accordingly. a recent proteomic and *O*-glycomic study has compared spontaneous or butyrate-induced Caco-2 differentiation and demonstrated that butyrate-stimulated differentiation was correlated with higher expression of sialylated *O*-glycan and lower fucosylation ^12^. Differentiated Caco-2 cells have also been applied to study intestinal epithelial interactions with biotherapeutics, microbiota by-products (e.g., short-chain fatty acids), or microbiota themselves more recently. Due to the fact that Caco-2 cells are easy to culture. it holds potential for use as a microbiome-based therapeutic assessment tool, attracting interest in characterizing and understanding the molecular alterations during its differentiation over time with a focus on host-microbiome interactions. In addition to the involvement in cancer development and anti-cancer therapy, lysine acetylation has also been demonstrated to be implicated in host-microbiome interactions in diseases, such as Crohn’s disease ^27^. However, to the best of our knowledge, there is no lysine acetylproteomic investigations when comparing spontaneous and DM-induced Caco-2 differentiations. Therefore, to better understand the Caco-2 cell differentiation processes with a particular focus on the host-microbiome interactions, this study applied proteomic, lysine acetyl-proteomic and glycoproteomic approaches to directly compare three commonly used Caco-2 differentiation protocols, including serum-containing and serum-free growth medium induced spontaneous differentiation and butyrate-containing DM medium induced rapid differentiation.

We showed that both spontaneous and DM-induced differentiated Caco-2 cells displayed overall proteomic, lysine acetyl-proteomic and glycoproteomic patterns moving away from those of undifferentiated cells. Over the course of the time series, the proteomic clustering patterns of the DM group exhibited notable differences with a more stable clustering pattern after 7 days, indicating that the differentiation process was either complete or minimally changing by 7 days, ultimately reaching a steady-state of the cells. This aligns with the timeline of corresponding published butyrate-induced differentiation methods ^8,12^. In contrast, both spontaneous differentiation groups (DFBS and SFM) continued to cluster further apart from the undifferentiated cells with increasing time up until the last 21-day time-point tested ^6,9,12^. These results suggest ongoing proteomic and post-translational changes associated with the progression of spontaneous differentiation take longer to reach their respective plateaus, or a more steady-state cell phenotype. This was also seen visually as by day 7 the DM group showed extensive dome formation whereas the DFBS and SFM groups were not as extensive at the same early time-point.

We also demonstrated through pathway analysis an increase in stimuli response proteins/modified proteins which are represented at the cell surface, in both cellular and extracellular membranes (vesicles, exosomes). This is reflected by increases in proteins with roles in membrane trafficking pathways and extracellular vesicles such as PIK3C3 and CD9. This reinforces that there are changes within the cell membrane as expected alongside the major morphological changes from Caco-2’s primary cancerous phenotype into an epithelial-like cell. Although a change in secreted vesicle quantity and size was not observed in preliminary investigations (data not shown), enriched pathways such as stimuli response, molecule binding, cell communication, and receptor activity suggest an increase in cell-to-cell cross-talk as well as increased interactions with the environment, such as chemicals and microbes, as the cells differentiate. This likely ties into the vital symbiotic relationship between human intestinal processes and the gut microbiome which reside within it ^29^. Glycoproteomics data also highlighted an increase in biological process involved in symbiotic interaction, and our study observed a significant increase in proteins and/or their altered PTM levels that are known to interact with and play key roles in host-microbiome homeostasis such as multiple galectins (Gal-1, Gal-2, Gal-3, Gal-4, and Gal-7) and folate-receptor alpha subunit (Supplementary Table 2-4)3.

An advantage of quantitative proteomics is to quantify multiple proteins, such as biomarkers, in a non-targeted manner. Accordingly, this study was able to quantify most known biomarkers of Caco-2 differentiation into intestinal epithelium-like cells, including ALP-intestinal type, MUC13, DPP4, TST, BHD2, and TAGLN, which all followed expected trends ^9,17–19^. Intestinal type ALP is the most well-known biomarker for intestinal cellular differentiation ^14,15^. We showed that the protein abundance of intestinal type ALP gradually increased along with differentiation, with the DM group peaked at day 7, while SFM and DFBS groups peaked at day 14 and 21, respectively (Fig. 2B). This trend was in agreement with the overall proteome pattern as well as the microscopic examination, indicating the rapid and efficient differentiation of Caco-2 cells using DM medium. Interestingly, in addition to the intestinal type ALP, we as well as a previous study both found that the two other types of ALPs (i.e., placental type and germ cell type) were also significantly upregulated upon DM-induced differentiation at as early as day 7, but not in spontaneous differentiations (Fig. 2 C-D) ^16^. These three types of ALPs may all contribute to the increased alkaline phosphatase activity in differentiated Caco-2 cells as generally measured using *p*-nitrophenyl phosphate (pNPP)-based colorimetric assay. In addition to ALPs, DM-induced differentiation also displayed higher abundance of MUC-13 at day 7 as well as day 14 and 21.

Similarly, villin proteins, the key components of intestinal villi structure, was found to be significantly up-regulated in the DM group at day 7, but not in DFBS and SFM groups at day 7 or 21 ^20,21^. Altogether, these findings suggest that full and efficient Caco-2 differentiation into a epithelial-like intestinal cells can be rapidly (within 7 days) achieved with butyrate-containing commercially available DM medium. Protein acetylation plays important roles in regulating translation, transcription, metabolism and immunity in both eukaryotes and bacteria and has been implicated in multiple cancers, making it a molecular mechanism of interest in characterizing the Caco-2 derived intestinal model ^27^.

Unlike proteomics which showed more uniform changes across differently differentiated groups, it was found that there was only 53 and 125 significantly changed acetylation sites in DM group at day 7 and 21, respectively, which is quite low compared to the 415 differentially abundant sites in SFM group and 377 sites in DFBS group at day 21. This mild change of Kac sites in DM group may be due to the presence of butyrate in the DM medium, as butyrate is a known histone deacetylase (HDAC) inhibitor ^31^. We found that, along with the Caco-2 cell differentiation across groups, most Kac sites on histone proteins (the most abundant Kac proteins) were down-regulated, indicating a decreasing trend of overall lysine acetylation level in differentiated cells compared to cancerous cells. Accordingly, more down-regulated Kac sites than up-regulated Kac sites were observed in spontaneous differentiation groups (SFM and DFBS) (Fig. 7A). The reduced histone protein Kac levels in differentiated Caco-2 cells, which favor the closed chromatin conformation, coincides with the proteomic observations showing down-regulated translation and proliferation related protein expressions.

The HDAC inhibitor butyrate in DM medium reversed this trend of overall Kac levels by increasing some Kac proteins, in particular at day 7 when more up-regulated Kac sites than down-regulated were obtained. These overall Kac protein changes were also reflected on the PCA score plot where the D7_DM and D21_DM groups clustered between the undifferentiated and spontaneous differentiation groups (D21_SFM and D21_DFBS). Of the Kac sites that are up-regulated in differentiated Caco-2 cells identified in this study, they were found to be enriched in mitochondria, stress response related proteins (such as HSP90 and ARP2), and ion transport or binding functions (such as CLIC1, annexins and S100 family proteins). These findings indicate that protein acetylation may be involved in regulating the intracellular homeostasis as well as the cell-cell interactions during differentiation. It is known that calcium (Ca^2+^) plays an important role in maintaining intestinal homeostasis, including signalling and transport which are key processes in nutrient sensing, transport processes, and endocrine regulation ^32–37^. Proteomic data in this study also showed an upregulation of calcium ion binding functions as well as cadherin binding functions (which require calcium ions to function in cell adhesion/junction formation) reinforcing the importance of calcium within intestinal-like cells. The S100 protein family of Ca^2+^ binding EF-hand proteins play roles in both intracellular and extracellular regulation ^22,38^. These include functions in differentiation, Ca^2+^ homeostasis, metabolism, and immune response ^38^. This study demonstrated an association between Caco-2 differentiation and the up-regulated expression of multiple S100 family proteins. This suggests a change in calcium signalling within Caco-2 cells during differentiation, with S100 proteins playing a role in the altered signalling pathways. Additionally, some annexins which are a family of non-EF-hand Ca^2+^ binding proteins, showed a similar association with Caco-2 differentiation. Annexins can form complexes and interact with S100 protein family proteins to play roles in membrane and vesicle trafficking and organization ^23,39^. We observed enrichment of some annexins, changes in multiple Kac sites on several annexin proteins (A2, S4, A5, A11), as well as a functional enrichment of membrane and vesicle/exosome components for both up-regulated proteins and Kac sites. Accordingly, cancerous cells (including colorectal cancer cells) display altered calcium signalling, which has become a target in cancer therapeutics, as calcium signalling regulates cell division, motility, and death with key roles in cancer cell migration and invasion ^40,41^. While the relationships may be complicated, the findings in this study may indicate a shift in calcium signalling during Caco-2 differentiation and protein lysine acetylation may regulate this molecular process and influence the dynamic cellular changes occurring during Caco-2 cell differentiation.

In summary, this study offers a comprehensive characterization for Caco-2 cell differentiation using proteomics, acetyl-proteomics, and glycoproteomics. We confirmed expected protein biomarker trends, and demonstrated an increase of proteins with functions in cell adhesion, response to stimuli, cell signalling, epithelium development, as well as the extracellular vesicles/space. This study also provided crucial insights into the roles of lysine acetylation in Caco-2 differentiation and demonstrated an up-regulation of lysine acetylated proteins related to mitochondria functions. These findings reflect a shift of differentiated Caco-2 cells towards a more interconnected epithelial-like cell network and the development of cellular functionalities commonly associated with host-microbiome interactions. Compared to spontaneous differentiation, full and efficient differentiation can be achieved within 7 days using butyrate-containing differentiation medium for culturing the cells. Following applications for the assessment of microbiomes from patients and healthy individuals, actionable molecular biomarkers can be identified that are indicative of benefits and/or risks of microbiome products on the host. Altogether this study improved our understanding of the Caco-2 differentiation processes, which will aid its application in biomarker and drug target discovery, the study of host-microbiome interaction, as well as assay development for quality assessment of GI tract-directed therapeutics moving forward.

## Materials and Methods

### Caco-2 cell culture and differentiation

Human colorectal adenocarcinoma (Caco-2) cells were obtained from the American Type Culture Collection and routinely maintained in Dulbecco’s modified Eagle’s medium (DMEM) with 10% heat-inactivated (HI) FBS at 37°C in a humidified 5% CO_2_ incubator. Cells from passages 30-48 were used in this study. Cells were all confirmed to be mycoplasma negative using a PCR mycoplasma detection kit (Thermo Fisher Chemicals, J66117) following manufacturers instructions.

For differentiation, Caco-2 cells were seeded (day 0) at a density of ∼1e^6^cells/cm^2^ in DMEM with 10% HI-FBS. On day 1, cells were switched into either serum-free DMEM (SFM) or DMEM with 10% HI-FBS (DFBS) or Corning® Intestinal Epithelium Differentiation Medium (Corning product number 355357) (DM). The Caco-2 cells were then cultured for 21 days in one of the three growth/differentiation medias, with media changes performed every other day. Cell imaging was performed using EVOS cell imaging system (Thermo Fisher Scientific). Cells were harvested on day 1, 3, 7, 14, and 21. Cells were harvested by removing culture medium, washing adherent cells twice with DPBS, and lifted with 0.25% Trypsin-EDTA. To inactivate the trypsin, medium with 10% HI-FBS was added at a ratio of 1:1 and the cells were then pelleted at 400g for 5min. Supernatant was discarded, and the cell pellet was washed with DPBS twice pelleting cells again at 400g for 5min. The supernatant was removed, and the cell pellets were stored at - 20°C until processing.

### Protein extraction, digestion and TMT labeling

The cells were lysed by resuspending frozen cell pellets in 200uL of lysis buffer (1% SDS in 100mM TEAB) and sonicating them with a QSonica Q700 water-chilled cup-horn sonicator at 50% amplitude, 10 seconds pulse on/off cycle, for 10 minutes active sonication time at 8°C. The lysate was then centrifuged at 16,000g for 10min at 4°C. Protein-containing supernatant was transferred to a new tube and protein concentrations was determined using the Pierce BCA Protein Assay Kit (Thermo Fisher Scientific, Cat #23225) following the manufacturer protocol. 150ug of protein lysates of each sample were reduced with 15 mM tris(2-carboxyethyl)phosphine (TCEP) at 55°C for 1 hour and alkylated with 25 mM iodoacetamide for 30 minutes at 21°C protected from light. Proteins were then precipitated by adding 6 volumes of ice-cold acetone at -20°C overnight. Samples were then centrifuged at 14,000g for 10 minutes at 4°C and the acetone was carefully decanted without dislodging the protein pellet. The pellet was then air dried for 5-10 minutes, resuspended with 75uL of 100mM TEAB containing 3mAU of Lysyl endopeptidase (1mAU LysC : 50ug protein), and incubated at 37°C in a thermomixer at 500rpm in the dark for 3 hours. 75uL of 100mM TEAB containing 3ug of MS-grade trypsin (1ug trypsin : 50ug protein) was then added and incubated overnight at 37°C in a thermomixer. The digestion was stopped by acidifying each sample with 35uL of 10% formic acid and the digests were then dried on a centrivap for further TMT labeling.

TMT labeling was performed according to our previously established dry TMT labeling workflow using an 11-plex TMT Mass Tagging Kit (Thermo Fisher Scientific, lot XB323735) ^13^. Briefly, dried peptide (150ug) was resuspended in 37.5uL of 100mM TEAB/20%ACN and 50ug of each sample was combined to generate a pool sample. The remaining 100ug peptides of each sample was added to 200ug of lyophilized TMT reagents ^13^. Samples were randomized into 6 TMT 11plex experiments with the 1^st^ and 11^th^ channel of each experiment as reference channels. The TMT reaction was performed by incubation at 25°C for 2 hours with shaking at 500rpm, and quenched with 4uL of 5% hydroxylamine for 15min at room temperature. The 11 samples of each TMT experiment were then combined (1100ug per mixture) and dried with a centrivap. 100ug of the mixture was fractionated using a high pH reverse phase fractionation kit (Thermo Fisher Scientific, Cat #84868) for proteomics, and the remaining 1mg peptides of each mixture were used for lysine acetyl-peptide and glycopeptide enrichment.

### Immunoaffinity enrichment of lysine acetylated peptides

The enrichment of lysine acetylated peptides was performed using a PTMScan HS Acetyl-Lysine Motif [Ac-K] Kit (Cell Signaling Technology, Cat #50071) following the manufacturer’s instructions. Briefly, approximately 1mg TMT-labelled peptides of each mixture were re-suspended in 1.5 mL of HS IAP bind buffer added to PBS washed magnetic Ac-K motif-antibody beads and incubated on a rotator for two hours at 4°C for peptide binding. Unbound peptides were then washed with IAP wash buffer and water, and the Kac peptides were then eluted with 50µl 0.15% TFA twice. Both elutes were combined and dried with a centrivap followed by desalting using Pierce C18 spin tips (Cat #84850). For desalting, the C18 columns were activated by washing three times with 50µL 100% acetonitrile (ACN) and equilibrated with 50µL 0.1% formic acid (FA); the sample was then re-suspended in 0.1% FA, added to the column, and followed by two more washes with 0.1% FA and elution with 80%ACN/0.1%FA. The eluted peptide was then dried using a centrivap prior to LC-MSMS analysis.

### Hydrophilic interaction liquid interaction chromatography (HILIC) for glycopeptides

Hydrophilic interaction solid phase extraction (HILIC SPE) was conducted as previously described with minor changes ^42^. TMT-labelled peptides of each mixture were solubilized in 50 µL 80 % acetonitrile (ACN) with 1% of trifluoroacetic acid (TFA) and loaded onto the microcolumn packed with 5 mg of polyhydroxylethyl beads (100 Å, 5 µm, PolyLC, Columbus, MA). After three washes with 50 uL of the same loading buffer, glycopeptides retained on SPE column were eluted by 50 µL 30 % of ACN with 0.1 % TFA. Eluted glycopeptides were lyophilized and then deglycosylated by incubation with 50 µL of 1 unit per microliter of PGNase F (New England Biolabs, Ipswich, MA) in 50 mM Tris-HCl (pH=7.5) overnight at 37 °C. Deglycosylated peptides were dried with Speed Vac and the samples were stored at -80 °C until MS analysis.

### HPLC-MS/MS analysis

Unenriched and fractionated peptides were analyzed using Orbitrap Fusion Lumos mass spectrometer (Thermo Fisher Scientific) coupled to an Easy-nLC 1200 system for UPLC (Thermo Fisher Scientific). The instrument was calibrated by infusion prior to analysis with Pierce FlexMix Calibration Solution (Cat #A39239). For each fraction, 20 ul 0.1% FA was used to resuspend peptides and 2 µl were analyzed by loading onto a NanoViper Acclaim pepmap 100 trap column (75 µm diameter, 20 mm length with 3 µm C18 beads) and desalting with 0.1% formic acid in water (solvent A), before separating on an NanoViper Acclaim pepmap C18 reverse-phase analytical column (50 µm diameter, 150 mm length with 2 µm C18 beads).

Chromatographic separation was achieved at a flow rate of 0.300 µl/min over 100 min in five linear steps as follows (solvent A was 0.1% formic acid in water, solvent B was 0.1% formic acid in 80% acetonitrile): initial, 2% B; 0 min, 25% B; 80 min, 40% B: 90 min, 95% B; 95 min, 95% B; 100 min. The eluting peptides were analyzed in data-dependent mode MSMS. A MS survey scan of 400−1600 m/z was performed in the Orbitrap at a resolution of 120,000, auto maximum injection time and the standard preset AGC target. The top speed mode was used to select ions for MS2 analysis with dynamic exclusion 20 s with a ± 10 ppm window. During the MS2 analyses precursors were isolated using a width of 0.7 m/z and fragmented by HCD followed by Orbitrap analysis at a resolution of 50,000. HCD precursors were fragmented with collision energy of 38% with auto maximum injection time and an AGC target of 250%.

Enriched Kac peptides were analysed using Orbitrap Exploris 480 mass spectrometer (Thermo Fisher Scientific) coupled with an UltiMate 3000 RSLCnano system (Thermo Fisher Scientific). Peptides were loaded onto a tip column (75 μm diameter, 150 mm length) packed with reverse phase beads (3 μm/120 Å ReproSil-Pur C18 resin, Dr. Maisch HPLC GmbH). A 60 min gradient of 5 to 35% (v/v) from buffer A (0.1% (v/v) formic acid) to B (0.1% (v/v) formic acid with 80% (v/v) acetonitrile) at a flow rate of 300 μL/min was used. The MS method consisted of a full MS scan from 350 to 1200 m/z at resolution 120,000, followed by data-dependent MS/MS scans with a first mass of 110 m/z, MS2 resolution of 45,000, isolation window of 0.7 m/z, and HCD collision energy of 36%. A dynamic exclusion duration was set to 60 s.

Enriched glycopeptides were analyzed using Orbitrap Exploris 480 mass spectrometer coupled with an Ultimate 3000 UHPLC. Peptides was were loaded via an Acclaim PepMap 100 trap column (C18, 5 μm, 100Å; Thermo Scientific, San Jose, CA) onto a Waters BEH C18 column (100 µm × 100 mm) packed with reverse phase beads (1.7 µm, 120-Å pore size, Waters, Milford, MA). A 45 minutes gradient from 5–30 % acetonitrile (v/v) containing 0.1 % formic acid (v/v) was performed at an eluent flow rate of 500 nL/min. Data-dependent acquisition with a cycle time of 1s was used. The MS method consisted of a full ms1 scan (resolution: 60, 000; AGC target: 300%; maximum IT: 50 ms; scan range: 350-1600 m/z) preceded by subsequent ms2 scans (resolution: 30,000; AGC target: 200%; maximum IT: 105 ms; isolation window: 1.3 m/z; first mass: 110m/z; NCE: 35%). To minimize repeated sequencing 1 of the same peptides, the dynamic exclusion was set to 30 sec and the ‘exclude isotopes’ option was activated.

### Bioinformatic data processing and statistical data analysis

Raw MS data for TMT-based proteomics were searched against a Homo Sapiens (human) reviewed (Swiss-Prot) canonical protein database from UniProtKB (downloaded on 2023/11/09, with 20,428 protein entries) using MaxQuant (v2.4.10.0) ^43,44^. Default parameters were used for MaxQuant database search with reporter ion MS2 mode. The TMT reagent lot-specific reporter ion isotopic distributions was used as isotope correction factors for each set of TMT labeling data. The quantification data was then summarized and normalized using MSstatsTMT ^45^ with the reference channels as bridge channel for different TMT experiments to generate quantitative protein group data matrix for further statistical analysis.

Lysine acetyl-proteomics data were processed using MaxQuant (v2.4.0.0) ^43,44^ with the same UniProtKB human database (downloaded on 2023/11/09, 20,428 protein entries). MaxQuant parameters were set the same as the proteomics data as described above, with the following exceptions: lysine acetylation (Acetyl (K)) was added as additional variable modification and the maximum missing cleavage sites was set as 3. MSstatsPTM ^46^ was then used to summarize and normalize the Kac site abundance data using the reference channels as bridge channel for different TMT experiments. To calculate the relative lysine acetylation levels on proteins, the proteomic data was used to adjust the Kac site abundance using MSstatsPTM as well. The proteome adjusted abundance of all identified Kac sites in all groups were compared using a pairwise t-test with FDR correction. The sites with FDR ≤ 0.05 and fold change (FC) ≥ 2 were deemed as significantly changed in this study.

Glycoproteomic data was processed by MSFragger (version 3.7) implemented in Fragpipe 18.0 with an UniProtKB human database (downloaded on 2023/02/15, 20,376 protein entries). Decoy and contaminants lists were added by FragPipe. A default TMT10 workflow was used with changing TMT10 to TMT11 in TMT interrogator tab. Two missed cleavages were allowed with +57.02146 on cysteine as fixed modification. +15.9949 on methionine and 0.9840 on asparagine were set as variable. Mass tolerance was set to 20 ppm for both precursor and fragment ions. Percolator was run with 1% false discovery rate (FDR) filter and TMT-integrator filtered peptide spectral match and peptide with minimum probability of 0.9 for quantification. All identified glycopeptides with N-X-S/T motif were then selected for further statistical data analysis.

Principal Component Analysis (PCA) was performed using all quantified proteins, Kac sites or glycoproteins with no missing values ^47^. For partial least squares – discriminant analysis (PLS-DA), all summarized and normalized data was uploaded to MetaboAnalyst ^48^ for data filtering to keep variables with valid values in ≥50% samples and data imputation using K-Nearest Neighbors Algorithm; the resulting data matrix was then used for PLS-DA analysis to obtain variable importance measures of each protein, Kac site or glycoprotein to the model. A PLS-DA model with a goodness of prediction (Q2) >0.4 was considered as acceptable in this study.

Hierarchical clustered was perfor med using Perseus ^49^ and R package *ComplexHeatmap*. Gene Ontology enrichment analyses were performed using STRING with all identified proteins as the background for the calculation of false discovery rates (FDR) ^50^.

### Data visualization

Experimental workflow was generated using BioRender (https://www.biorender.com/). Protein-protein interaction network was generated using STRING (https://string-db.org/). Boxplots, volcano plots and functional enrichment dot plots were generated using R package *ggplot2* and *ggpubr*. Heatmap, hierarchical clustering, and UpSet analysis and plotting were performed using R package *ComplexHeatmap*. PCA and plotting was performed in R as well using R function *autoplot* and *prcomp*.

### Data availability

All MS proteomics data that support the findings of this study have been deposited to the ProteomeXchange Consortium (http://www.proteomexchange.org).

## Acknowledgement

This work was supported by the Government of Canada through Health Canada. We gratefully acknowledge Dr. Simon Sauvé, Dr. Michael Rosu-Myles, Dr. Xuguang Li and Dr. Roger Y. Tam from Health Canada for commenting on the manuscript.

