## Supplementary materials for "Multilevel proteomic profiling of colorectal adenocarcinoma cell differentiation to characterize an intestinal epithelial model"

*^1^Regulatory Research Division, Biologic and Radiopharmaceutical Drugs Directorate, Health Products and Food Branch, Health Canada, Ottawa, Canada; ^2^Department of Biochemistry, Microbiology and Immunology, Faculty of Medicine, University of Ottawa, Ottawa, Canada; ^3^School of Pharmaceutical Sciences, Faculty of Medicine, University of Ottawa, Ottawa, Canada; ^4^Human Health Therapeutics Research Centre, National Research Council Canada, Ottawa, Ontario, Canada.*

*Correspondance: X.Z.,**; R.C.,*

**Supplementary Tables**

Supplemental Table 1. Outline of sample distribution across TMT experiments and labels used.

| **Channels** | **Mixture 1** | **Mixture 2** | **Mixture 3** | **Mixture 4** | **Mixture 5** | **Mixture 6** |
| --- | --- | --- | --- | --- | --- | --- |
| TMT^10^-126 | Pool | Pool | Pool | Pool | Pool | Pool |
| TMT^10^-127N | D7-SFM-1 | D3-DM-1 | D3-DFBS-1 | D21-SFM-4 | D14-SFM-2 | D14-DM-4 |
| TMT^10^-127C | D1-undiff-3 | D7-DFBS-1 | D7-SFM-3 | D14-DFBS-2 | D7-SFM-4 | D3-SFM-4 |
| TMT^10^-128N | D14-DFBS-1 | D1-undiff-2 | D14-DM-1 | D3-SFM-2 | D3-DM-3 | D7-DM-1 |
| TMT^10^-128C | D21-DFBS-1 | D3-SFM-1 | D21-DFBS-2 | D21-DM-2 | D7-DFBS-3 | D21-SFM-3 |
| TMT^10^-129N | D3-DFBS-3 | D21-DM-3 | D7-DM-3 | D14-SFM-3 | D14-DFBS-4 | D7-SFM-2 |
| TMT^10^-129C | D7-DM-2 | D1-undiff-5 | D3-DM-4 | D3-DFBS-2 | D1-undiff-4 | D21-DFBS-4 |
| TMT^10^-130N | D21-SFM-2 | D21-DFBS-3 | D14-DFBS-3 | D1-undiff-1 | D14-DM-2 | D14-SFM-1 |
| TMT^10^-130C | D14-DM-3 | D3-DFBS-4 | D21-SFM-1 | D7-DM-4 | D21-DM-1 | D7-DFBS-2 |
| TMT^10^-131N | D21-DM-4 | D14-SFM-4 | D7-DFBS-4 | D1-undiff-6 | D3-SFM-3 | D3-DM-2 |
| TMT^11^-131C | Pool | Pool | Pool | Pool | Pool | Pool |

Supplemental Table 2. Compilation of proteomic data, including PLSDA VIP, Log2FC and adjusted p-value of key differentiation groups compared to undifferentiated cells, Log2(Normalized Intensity) of all replicates for compared groups. **Excel Sheet*

Supplemental Table 3. Compilation of glycomic data, including PLSDA VIP score and Log2(Normalized Intensity) of key groups. **Excel Sheet*

Supplemental Table 4. Compilation of acetylomic data, including PLSDA VIP, Log2FC and adjusted p-value of key differentiation groups compared to undifferentiated cells, Log2(Normalized Intensity) of all replicates for compared groups. **Excel Sheet*

**Supplementary Figures**

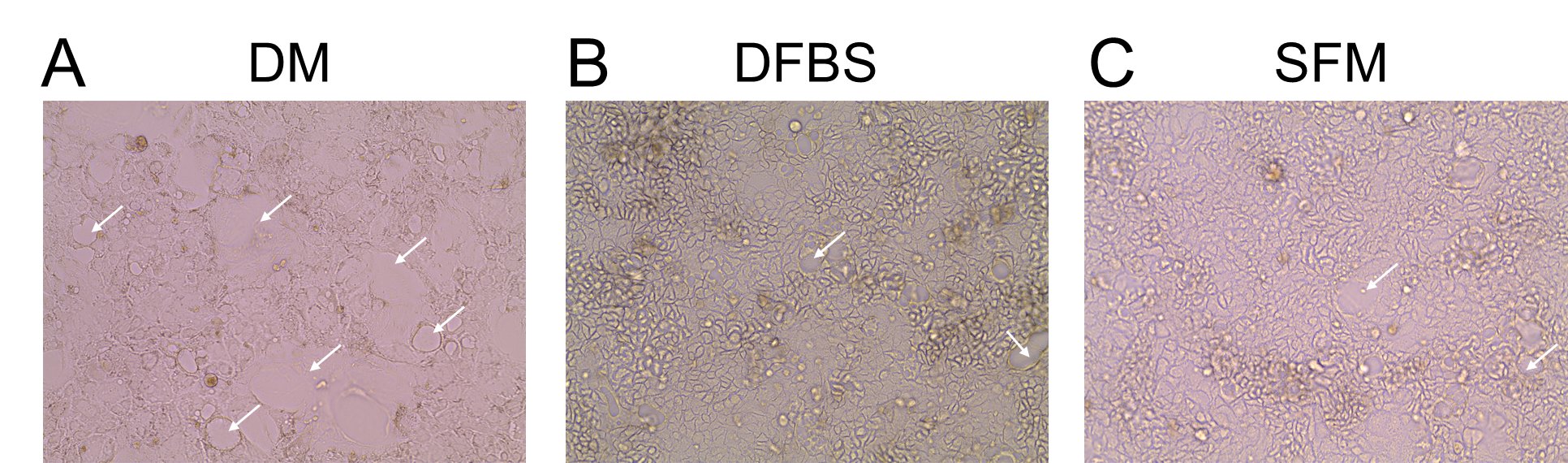

*Supplemental Figure 1. Microscopic images comparing cell morphology during the differentiation of Caco-2 cells. EVOS bright-field image of Caco-2 cells cultured for 7 days in DM (A), DFBS (B), SFM (C). White arrows indicate dome-like structure, an indicator of intestinal cellular differentiation.*

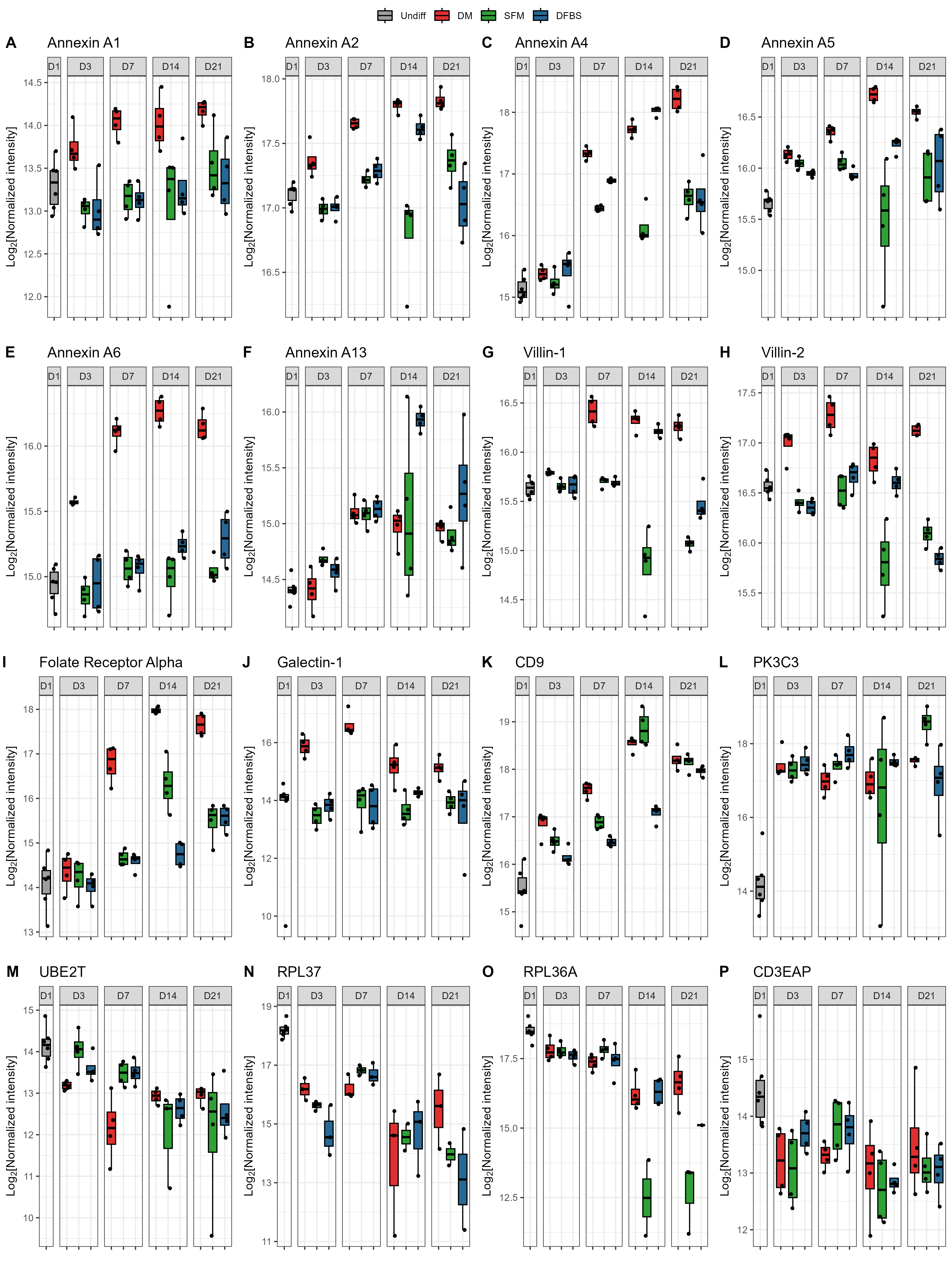

*Supplemental Figure 2. Differential protein expression of selected proteins in key Caco-2 cell differentiation groups. Log_2_(Normalized Intensity) of Annexin A1 (A), Annexin A2 (B), Annexin A4 (C), Annexin A5 (D), Annexin A6 (E), Annexin A13 (F), Villin-1 (G), Villin-2 (H), Folate Receptor Alpha (I), Galectin-1 (J), CD9 (K), PK3C3 (L), UBE2T (M), RPL37 (N), RPL36A (O), CD3EAP (P) over time.*

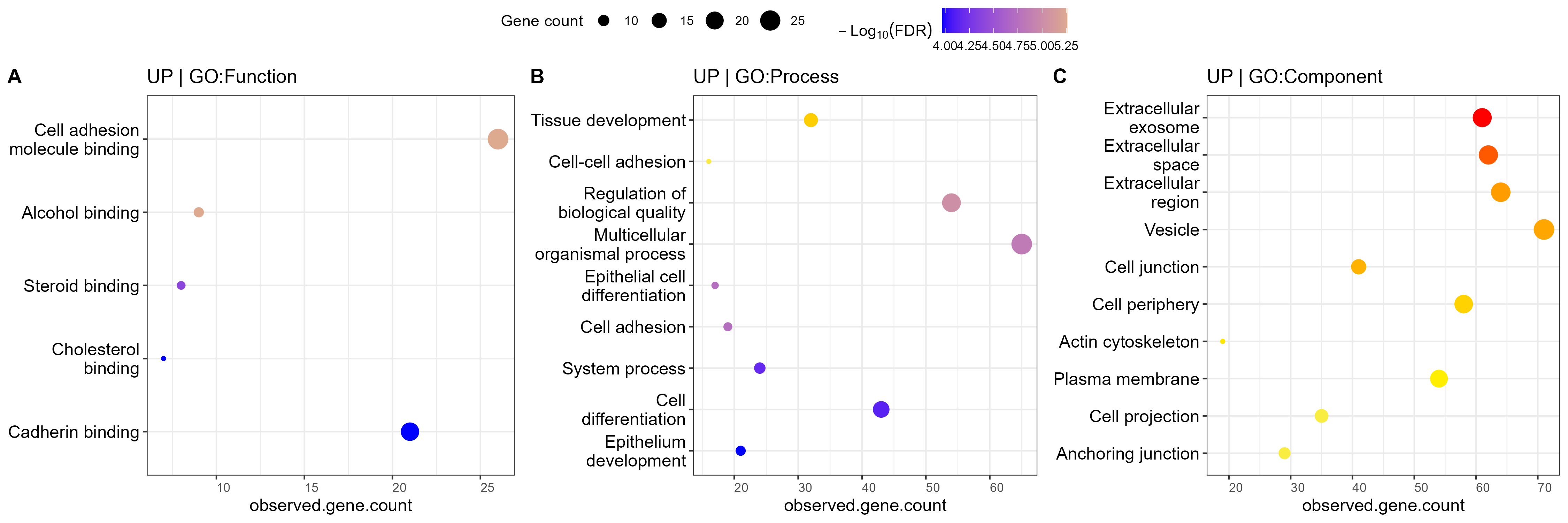

*Supplemental Figure 3. Functional enrichment analysis of proteins collected from differentiated and undifferentiated Caco-2 cells which were further upregulated in the D7DM group as compared to D21DFBS and D21SFM groups. Gene Ontology enrichment analysis using STRING of PLSDA VIP ≥ 1 proteins up-regulated functions (A), processes (B), and components (C).*

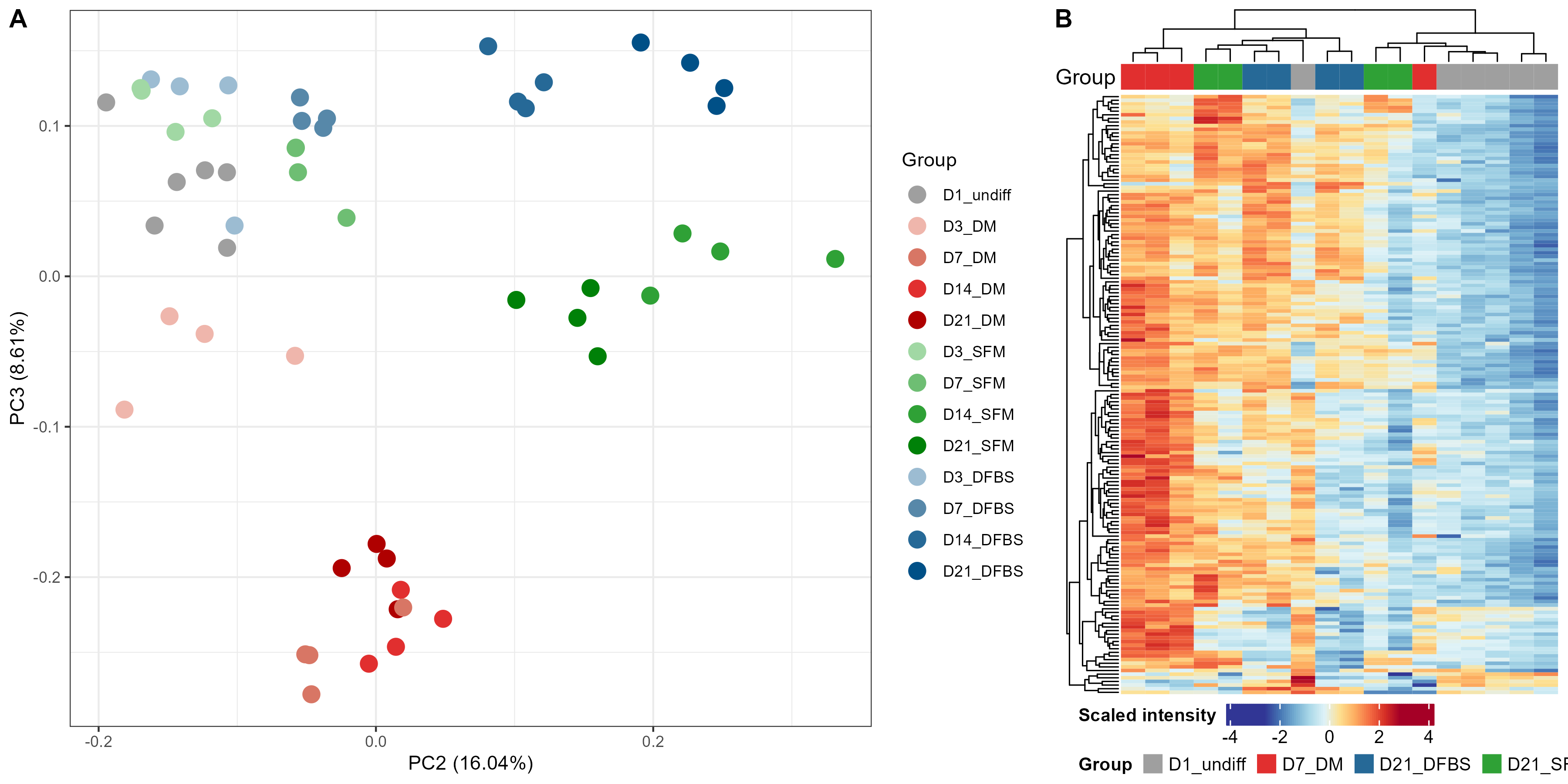

*Supplemental Figure 4. Overall trends in N-glycosylated proteins of Caco-2 differentiation from carcinoma into intestinal epithelial-like cells. PCA analysis (A). Heatmap and clustering of all Q100 N-glycosylated proteins (B).*

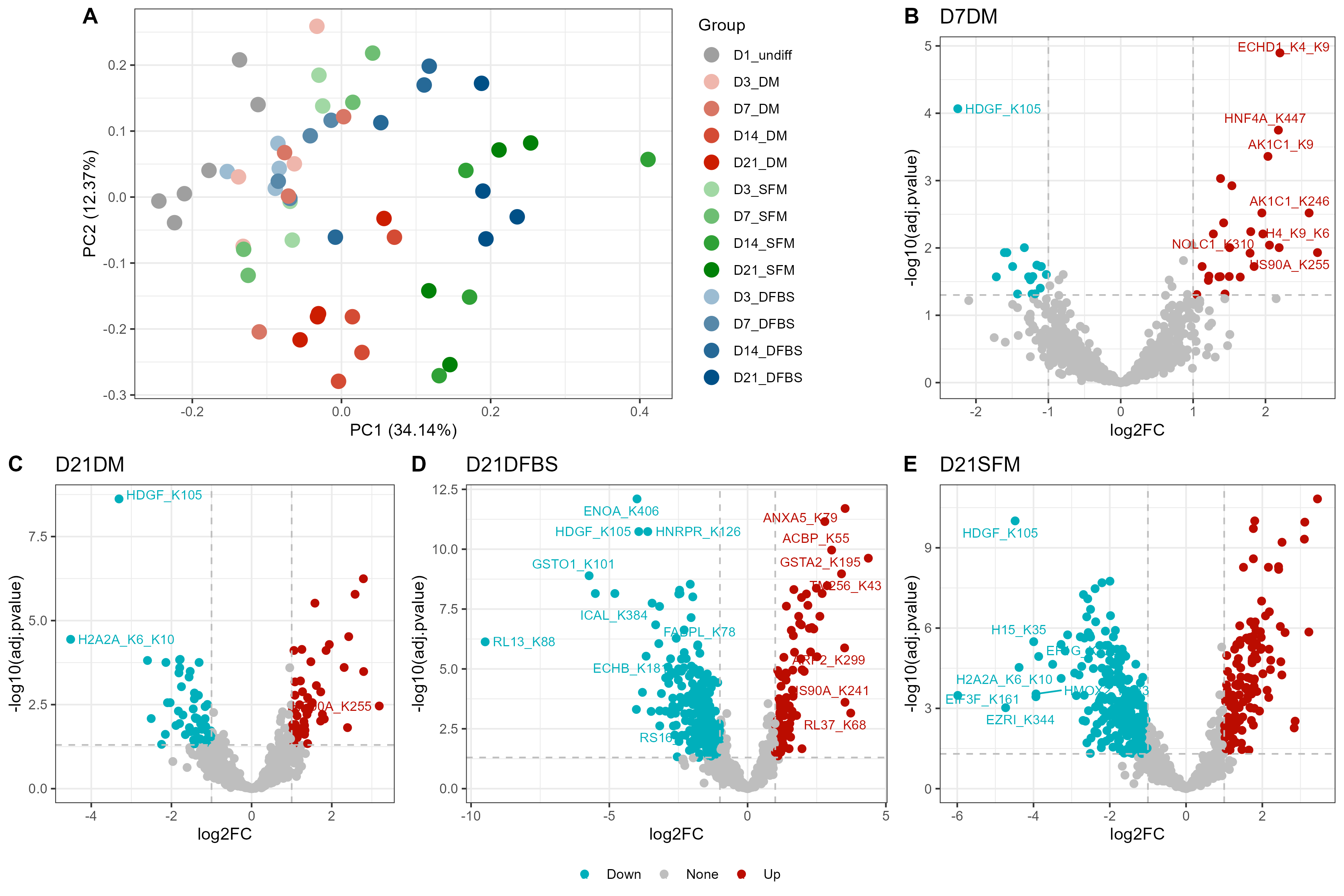

*Supplemental Figure 5. Overall trends in acetylated proteins of key differentiated Caco-2 cell groups compared to undifferentiated cells. PCA analysis (A). Volcano plot of Log2FC of all of all lysine acetylated proteins, highlighting those with a fold change ≥ 2 and adjusted p-value ≤ 0.05 comparing undifferentiated day 1 Caco-2 cells to cells after 7 days of growth in DM (B), or 21 days of growth in DM (C), DFBS (D), and SFM (E).*

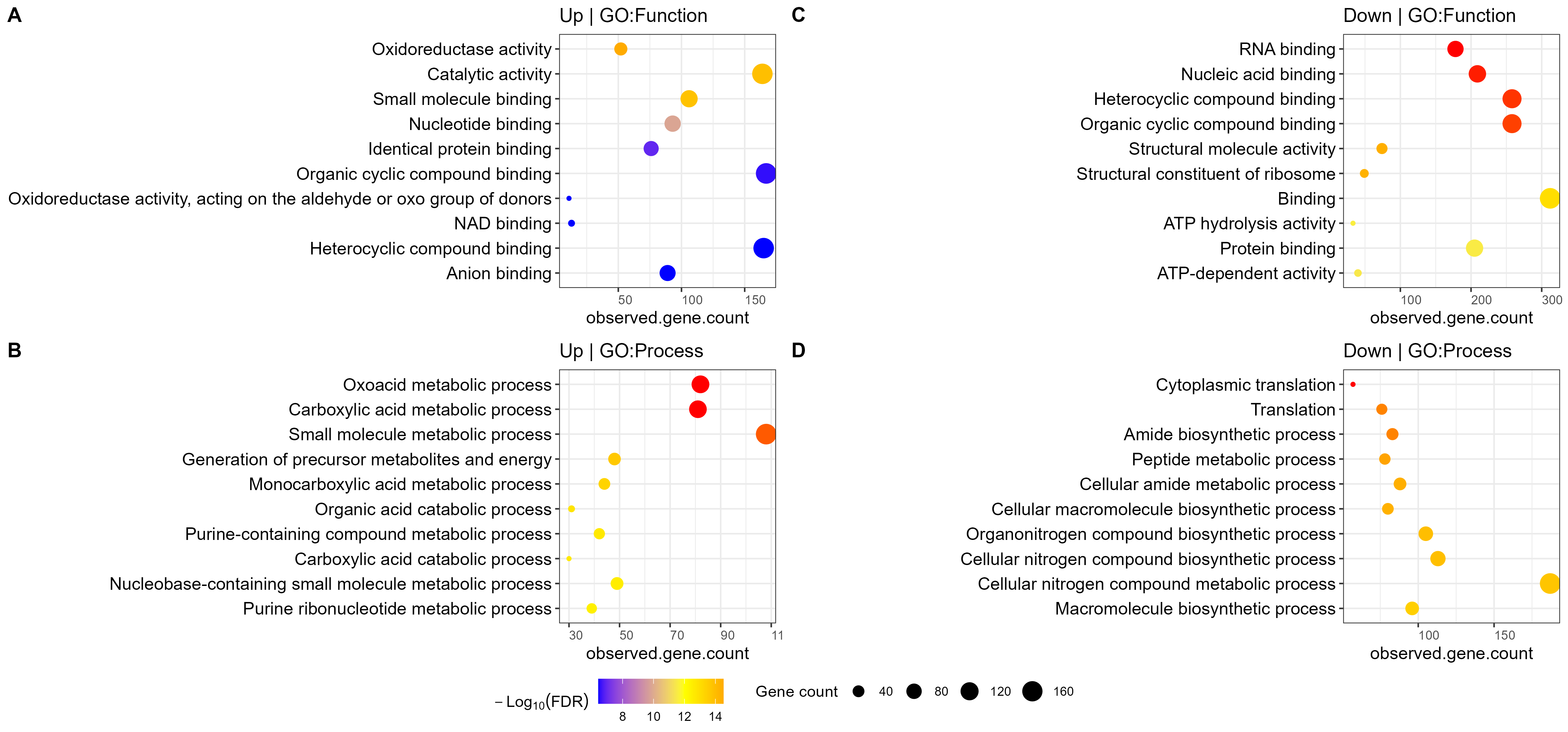

*Supplemental Figure 6. Functional enrichment and enriched protein network of K-acetylated proteins in differentiated cells (D7DM, D21DFBS, D21SFM) compared to undifferentiated (D1undiff) cells. Gene Ontology enrichment analysis using STRING of PLSDA VIP ≥ 1 proteins upregulated functions (A) and processes (B); down regulated functions (D) and processes (E).*
